# Weak effect of *Gypsy* retrotransposon bursts on *Sonneratia alba* salt stress gene expression

**DOI:** 10.1101/2021.03.24.436880

**Authors:** Yushuai Wang, Aimei Dai, Tian Tang

## Abstract

**Background and Aims:** Transposable elements (TEs) are an important source of genetic diversity and can be co-opted for the regulation of host genes. However, to what extent the pervasive TE colonization of plant genomes has contributed to stress adaptation remains controversial. Plants inhabiting harsh environments in nature provide a unique opportunity to answer this question.

**Methods:** We compared TE compositions and their evolutionary dynamics in the genomes of two mangrove species: the pioneer *Sonneratia alba* and its less salt-tolerant relative *S. caseolaris*. Age distribution, strength of purifying selection and the removal rate of LTR (long terminal repeat) retrotransposons were estimated. Phylogenetic analysis of LTR retrotransposons and their distribution in the genome of *S. alba* were surveyed. Small RNA sequencing and whole-genome bisulfite sequencing was conducted using leaves of *S. alba*. Expression pattern of LTR retrotransposons and their nearby genes were examined using RNA-seq data of *S. alba* under different salt treatments.

**Key Results:** *S. alba* possesses more TEs than *S. caseolaris*. Particularly, many more young *Gypsy* LTR retrotransposons have accumulated in *S. alba* than in *S. caseolaris* despite an increase in purifying selection against TE insertions. The top two most abundant *Gypsy* families in *S. alba* preferentially insert in gene-poor regions. They are under relaxed epigenetic repression, probably due to the presence of CHROMO domains in their 3’-ends. Although a considerable number of TEs in *S. alba* showed differential expression under salt stress, only four copies were significantly correlated with their nearby genes in expression levels. One such TE-gene pair involves *Abscisic acid 8’-hydroxylase 3* functioning in abscisic acid catabolism.

**Conclusions:** This study sheds light on the evolutionary dynamics and potential function of TEs in an extremophile. Our results suggest that the conclusion on co-option of TEs should be cautious even though activation of TEs by stress might be prevalent.

## INTRODUCTION

Transposable elements (TEs) are genetically diverse mobile sequences that can move to new sites in genomes by a “copy-and-paste” mechanism via an RNA intermediate (Class I, retrotransposons) or a “cut-and-paste” mechanism (Class II, transposons) (Wicker et al., 2007, Bourque et al., 2018). TE insertions often disrupt gene function or regulation and induce ectopic/nonallelic recombination events (Chuong et al., 2017). As a result, their activities are repressed by host epigenetic silencing and small interfering RNAs (siRNAs) (Fultz et al., 2015). Bursts of TE abundance are further subject to mutational decay and are eliminated by genome purging mechanisms in the long run. In many higher eukaryotes TEs constitute more than half of the DNA (Fedoroff, 2012) and play a critical role in the evolution of genome size and organization (Lisch, 2013). It is thus important to dissect the evolutionary dynamics of TEs in the genome.

While TEs are often deleterious or effectively neutral, there is growing evidence that TE activity can help the host adapt to environmental challenges (Casacuberta and Gonzalez, 2013, Horvath et al., 2017). High copy number and mutagenicity make TEs a major source of genetic innovation that can be activated by environmental cues (Capy et al., 2000, Grandbastien, 2015, Rey et al., 2016). This may accelerate mutation rates and generate variability upon which natural selection could act. Indeed, recent findings show that TEs can facilitate rapid adaptation to novel environments in species with limited genetic variation, such as *Capsella rubella* the annual inbreeding forb (Niu et al., 2019) and *Cardiocondyla obscurior* the invasive inbreeding ant (Schrader et al., 2014). Meanwhile, some TEs are known to carry stress-responsive regulatory elements. These elements, once activated by stress, may have a large role in remodeling regulatory networks and reshaping transcriptomes (Naito et al., 2009, Cavrak et al., 2014, Grandbastien, 2015, Chuong et al., 2017). Despite many reports that individual TE integration events result in changes in gene expression or even phenotypes under stress conditions (Butelli et al., 2012, Lisch, 2013), there is empirical evidence that regulatory function of TEs is not always strong (de Souza et al., 2013, Simonti et al., 2017).

Mangroves are a group of phylogenetically diverse woody species that invaded and adapted to the extreme and extremely fluctuating environments in the tropical intertidal zone (Tomlinson, 1986). These species have undergone multiple population bottlenecks during their evolutionary history and typically harbor extremely low standing genetic variation (Zhou et al., 2011, Guo et al., 2018). A recent study reported massive and convergent TE load reduction in mangroves, when compared to non-mangrove species (Lyu et al., 2018). However, we found a continuous host-TE battle masked by the TE load reduction in *Rhizophora apiculata*, the tall-stilted mangrove, suggesting TE activities may enable mangroves to maintain genetic diversity (Wang et al., 2018). It still remains unclear how often TE activities are significant to the evolution of mangrove genomes and to what extent TEs may have influenced gene expression during mangrove stress adaptation.

In this study, we surveyed the evolutionary dynamics and potential regulatory functions of TEs in *Sonneratia alba* Sm., the pioneer mangrove (Tomlinson, 1986), in comparison with its relative *S. caseolaris* (L.) Engl.. Although both are mangroves, *S. alba* is more salt-tolerant than *S. caseolaris* and often grows on the seaward side of mangrove forests with maximal growth in 5 to 50% sea water (Ball and Pidsley, 1995). In contrast, *S.caseolaris* is common in the inner parts of mangrove forests (Bunt et al., 1982) and in some instances has also been found growing in fully fresh water (Rahim and Bakar, 2018). Here we report recent amplification of *Gypsy* LTR (long terminal repeat) retrotransposons in *S. alba.* Nevertheless, this burst of *Gypsy* retrotransposons has only a weak effect on salt stress gene expression. Our results suggest that the conclusion on co-option of TEs should be cautious even though activation of TEs by stress might be prevalent.

## MATERIALS AND METHODS

### Annotation and classification of transposable elements

Whole genome sequences of *Sonneratia alba*, *Sonneratia caseolaris*, and *Eucalyptus grandis* were retrieved from previous studies (Myburg et al., 2014, Lyu et al., 2018). LTR retrotransposons were *de novo* mined and classified using work flows described previously (Wang et al., 2018). The annotation of long interspersed nuclear elements (LINE), short interspersed nuclear elements (SINE), and DNA transposons (MULE, hAT, PIF, CACTA and TcMar) was carried out as follows. First, sequences of the most conserved coding domain of each TE family were retrieved form the NCBI database (Coordinators, 2018) and used as queries against each genome using the TARGeT pipeline (Han et al., 2009) with the *e*-value cutoff of 0.001. Then, the identified homologs together with their 5 kb up- and downstream sequences were aligned within individual TE families using MAFFT (Katoh et al., 2002) and resolved into lineages using phylogenetic analyses as implemented in SiLiX (Miele et al., 2011). The terminals of each putative element were determined manually according to sequence similarity with its closely related elements. Miniature inverted-repeat transposable elements (MITEs) and Helitron copies were identified using MITE_HUNTER (Han and Wessler, 2010) and HelitronScanner (Xiong et al., 2014) with default parameters, respectively. Finally, all *de novo* identified TEs were added to RepBase (v.23.06) (Bao et al., 2015) for each genome separately and used to annotate TE contents in each species using RepeatMasker (v.4.0.0) (Smit et al., 2013-2015).

### Dating LTR retrotransposon elements

Age of each LTR retrotransposon element was estimated based on sequence divergence between their LTR pair (SanMiguel et al., 1998). The pairs from individual LTR retrotransposons were aligned using MUSCLE (v.3.8.31) with default parameters (Edgar, 2004). Sequence divergence *k* between each pair of LTRs was estimated using Kimura’s two-parameter model (Kimura, 1980). The age (T) of each LTR retrotransposons was estimated using the formula T = *k* / 2*r*, where *r* is the substitution rate of 1.3 × 10^−8^ substitutions per site per year (Ma and Bennetzen, 2004).

### Distribution of TEs across genomic regions

The insertion frequency of TEs categories in intergenic regions, introns, and coding regions in *S. alba* and *S. caseolaris* were estimated according to the procedure described before (Wright et al., 2003). Briefly, the genome sequences and annotated gene positions of *S. alba* and *S. caseolaris* were obtained from Lyu et al. (2018). For each species, each scaffold (>100 kb) was submitted to a BLAST search against our TE database to determine the positions of each TE superfamily along the entire set of scaffolds. TE positions were then verified manually to eliminate all redundancies and record the presence of nested insertions. To estimate gene density, sequences were scanned using 100 kb sliding windows with 50 kb steps across each scaffold. Gene density of each window was calculated as the fraction of the total length occupied by coding sequences.

### Phylogenetic analyses of LTR retrotransposons

Multiple alignments of full-length sequences of intact *Copia* or *Gypsy* retrotransposons were conducted using MAFFT (Katoh et al., 2002). Maximum likelihood phylogenetic trees were inferred with FastTree (Price et al., 2009) using a generalized time-reversible (GTR) model of nucleotide substitution. The generated trees were visualized with Figtree (v.1.4.2) (Rambaut, 2012). Resampling of estimated log-likelihoods (RELL)-like local support values was performed using FastTree (Price et al., 2009) with default parameters.

### Identification and sequence analysis of the chromodomain in LTR retrotransposons

Chromodomains residing in LTR retrotransposons were identified by tBLASTx (NCBI) searches against the *nr* database (NCBI) with default parameters (Boratyn et al., 2012). Multiple alignments of the chromodomain amino acid sequences were performed using MAFFT (Katoh et al., 2002). Tertiary structures of the chromodomains peptides were predicted using the Phyre2 server with default parameters (Kelley et al., 2015). Known chromodomain amino acid sequences were retrieved from the NCBI database (Coordinators, 2018), including DmHP1 (GenBank accession number, M57574) from *Drosophila melanogaster*, MAGGY (L35053) and Pyret (AB062507) from *Magnaporthe grisea*, Cft1 (AF051915) from *Cladosporium fulvum*, PpatensLTR1 (XM_001752430) from *Physcomitrella patens*, Tf2-12 (RVW30894) from *Vitis vinifera*, sr35 (AC068924), rn8 (AK068625) and RIRE3 (AC119148) from *Oryza sativa,* Tekay (AF050455) from *Zea mays,* RetroSor2 (AF061282) from *Sorghum bicolor*, and Tma (AF147263) from *Arabidopsis thaliana*.

### Epigenome sequencing and analyses

Young leaves of *S. alba* were collected from Qinlan Harbour, Hainan, China. Genomic DNA and total RNA were extracted separately using the modified CTAB protocol (Yang et al., 2008). Bisulfite (BS-seq) and small RNA sequencing were conducted with two biological replicates each according to previously published procedures (Wen et al., 2016, Wang et al., 2018).

Bisulfite sequencing reads were filtered for quality using Trimmomatic (v.0.32) (Bolger et al., 2014) and mapped to the *S. alba* genome using Bismark (v.0.16.3) (Krueger and Andrews, 2011) with default parameters. Only uniquely mapping reads were retained. Methylation levels of each LTR retrotransposon copy were calculated as the proportion of Cs among Cs and Ts (#C/(#C+#T)) by methylation context (CG, CHG and CHH) using custom Perl scripts. Only sites covered by more than three reads were used for the calculation. Results were visualized using R (v.3.1.3) (https://www.R-project.org).

Small RNA sequencing reads were quality controlled following procedures published previously (Wen et al., 2016) while structural non-coding RNAs, such as ribosomal RNA (rRNA), transfer RNA (tRNA), small nuclear RNA (snRNA), small nucleolar RNA (snoRNA), and known microRNAs (miRNAs) were removed. The remaining reads were considered as putative siRNAs and aligned to *S. alba* genomes using Bowtie (Langmead et al., 2009) with no mismatches allowed. SiRNA density of a particular LTR retrotransposon element was calculated as read counts per kilo base pair per element. Multiply-mapping reads were weighted by the number of their mapping locations. All statistical calculations were performed in R (v.3.1.3).

### Expression analyses of TEs and TE-flanking genes under salt treatments

Transcriptome data from *S. alba* under the 0, 250mM, and 500mM NaCl treatments were obtained from Feng et al. (2020) (NCBI Sequence Read Archive database accession No. PRJNA615770). Sequencing adapters and low quality reads were filtered using Trimmomatic (v.0.32) (Bolger et al., 2014). Clean reads were mapped to coding genes or intact LTR retrotransposons separately using Bowtie (Langmead et al., 2009) allowing no more than two mismatches. We calculated the expression level of each LTR retrotransposon element as reads per kilobase of transcript. Expression levels of individual genes were analyzed using HTSeq (v.0.6.1) (Anders et al., 2015) with the parameter -s set to “no” and normalized to Reads Per Kilobase per Million mapped reads (RPKM). Only genes located within 3 kb up- or downstream of the insertion of an intact or truncated LTR retrotransposon were considered in our analysis. The expression differences of TEs or genes across salt treatments were identified using DESeq2 (Love et al., 2014) requiring Benjamini-Hochberg multiple testing corrected (Benjamini and Hochberg, 1995) FDR < 0.05 and ≥ 2-fold change.

### Identification of *cis*-acting regulatory elements (CREs) of LTR retrotransposons

The CREs within LTRs of the consensus sequence of each LTR retrotransposon family or within the whole sequence of each truncated LTR retrotransposon were identified using PlantCARE (http://bioinformatics.psb.ugent.be/webtools/plantcare/html/search_CARE.html) (Lescot et al., 2002). *Gypsy* and *Copia* families that have more than ten copies were considered in this analysis.

### MiRNA target prediction

Plant mature miRNAs and their precursors were retrieved from miRBase (Release 22) (Kozomara et al., 2019). The identification of the known and novel miRNAs was performed by miRDeep2 software package (Friedlander et al., 2012) with the default parameters. The expression level of each identified miRNAs was normalized to Reads Per Million mapped reads (RPM). MiRNAs that with a RPM > 1 in both biological replicates were considered expressed and retained for further analysis. The prediction of the putative target genes for the known and novel miRNAs was done with the psRNATarget (Dai and Zhao, 2011) using an expectation score of 2.5 as the cutoff in the prediction. All LTR retrotransposons including the intact and truncated elements and solo-LTRs were used in these analyses.

### Accession numbers

Sequencing data from this article can be found in the GenBank data libraries under accession number PRJNA636918.

## RESULTS

### TEs are more abundant in *S. alba* than in *S. caseolaris*

*S. alba* and *S. caseolaris* diverged about ∼11.6 million years (Myr) ago (Yang et al., 2016). The whole genome sequences of *S. caseolaris* (∼208Mb, 1*C* = 259 Mb) and *S. alba* (∼207Mb, 1*C* = 284 Mb) have recently been *de novo* assembled (Lyu et al., 2018). We used the whole genome of *Eucalyptus grandis* (Myburg et al., 2014) as the outgroup following a previous study (Xu et al., 2017) and studied the evolutionary dynamics of TEs in *S. alba* in comparison with *S. caseolaris.* We annotated TEs in each species considering both TE structure and sequence homology and clustered them into families using the same procedure (see *MATERIALS AND METHODS*).

Consistent with the report of TE load reduction in mangroves (Lyu et al., 2018), we found TEs only accounted for 5.3% (11,091,467 bp) and 11.0% (22,805,675 bp) of the *S. caseolaris* and *S. alba* genomes, respectively, far less than the proportion of TEs in the *E. grandis* genome (27.7%, 191,425,687 bp, Figure 1A). In all three species, LTR retrotransposons are the dominant TE components (Figure 1A). Nevertheless, *Gypsy* retrotransposons are more abundant than *Copia* retrotransposons in the two *Sonneratia* species while the opposite pattern is observed in the outgroup *E. grandis* (Figure 1A). The expansion of *Gypsy* retrotransposons is more prominent in *S. alba* (12.7 Mb or 6.1%) than in *S. caseolaris* (3.3 Mb or 1.6%, Figure 1B). We identified 947 intact LTR retrotransposon copies comprising 180 *Copia* and 767 *Gypsy* elements in *S. alba* and 125 intact LTR retrotransposon copies consisting of 51 *Copia* and 74 *Gypsy* elements in *S. caseolaris* (Supplementary Data: Table S1). Although mangroves have experienced an overall reduction in TE load (Lyu et al., 2018), more *Gypsy* retrotransposons have accumulated in *S. alba* than in *S. caseolaris*.

**Figure 1.**
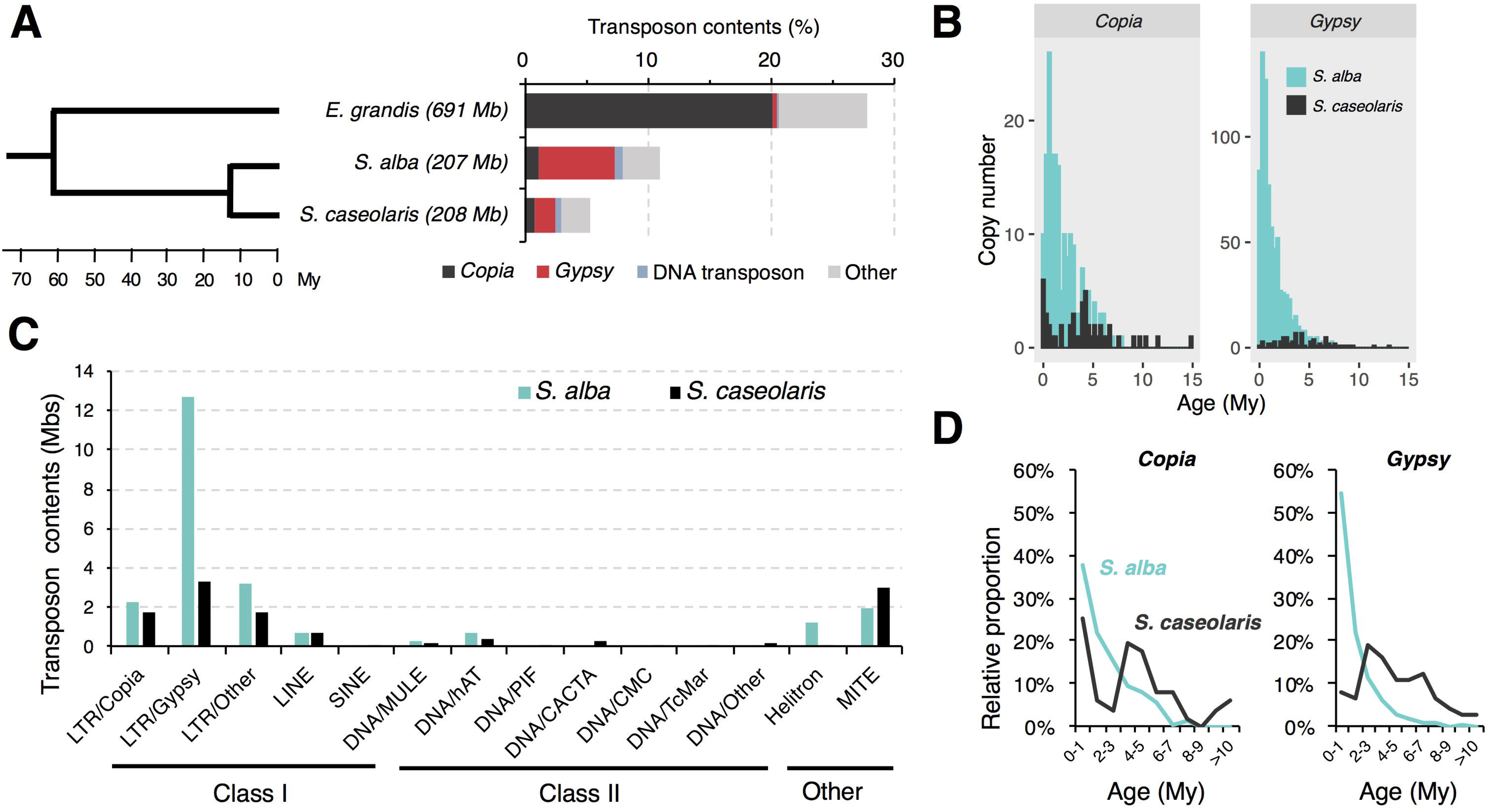
TE composition in *Sonneratia alba* and *S. caseolaris*. (A) Bar plot shows the proportions of transposon groups in *S. alba, S. caseolaris*, and *Eucalyptus grandis* (the outgroup). Tree topology is adopted from Lyu et al. (2018). (B) Histogram shows the distribution of transposon contents across families between *S. alba* and *S. caseolaris*. (C) Histogram shows the copy number of *Copia* and *Gypsy* elements in *S. alba* and *S. caseolaris* grouped by age. (D) Line graph shows the relative proportion of total *Copia* and *Gypsy* elements in different age bins.

### *Gypsy* retrotransposons experienced recent bursts in *S. alba*

To explore the dynamics of TE accumulation, we first estimated ages of the intact *Gypsy* and *Copia* retrotransposons by comparing LTR sequences flanking each element and assuming a molecular clock of 1.3×10^8^ (Ma and Bennetzen, 2004). We found these retrotransposons were much younger in *S. alba* (*Copia*: 1.47 Myr old; *Gypsy*: 0.87 Myr old) than in *S. caseolaris* (*Copia*: 3.85 Myr old and *Gypsy*: 3.99 Myr old). These differences are highly significant (Mann-Whitney *U*-test, both P < 0.001, Figure 1C). The recent burst of TE activity is most striking for *Gypsy* elements in *S. alba*. Young (< 1 Myr old) *Gypsy* elements account for 54.4% of the intact elements in *S. alba* but only 8.1% in *S. caseolaris* (Figure 1D). By contrast, the proportions of young *Copia* elements are more similar between *S. alba* (37.9%) and *S. caseolaris* (25.5%, Figure 1D). Therefore, there are more recent originations of the *Gypsy* element in *S. alba* than in *S. caseolaris*.

We then inferred the intensity of purifying selection against deleterious TE insertions by comparing TE distribution in relation to coding regions. We found TE insertions are strongly under-represented in coding regions compared to introns and intergenic regions (Table 1), consistent with the prediction of the deleterious insertion model (Finnegan, 1992). The abundance of all TE categories regardless of their locations are significantly lower in *S. alba* than in *S. caseolaris* (Table 1), suggesting TEs are under stronger purifying selection in the former species. However, the abundance of *Gypsy* elements in introns and intergenic regions was comparable between *S. alba* and *S. caseolaris* (Mann-Whitney *U*-test, all P > 0.05), suggesting that the insertions of *Gypsy* into introns and intergenic regions have experienced relaxation of purifying selection in *S. alba.* It is also possible that purifying selection against *Gypsy* insertions into coding regions is even stronger than that for other TE families in *S. alba*; otherwise, we should see more *Gypsy* insertions in the coding regions of that species.

**Table 1.** Location of TE insertions in relation to coding regions in *S. alba* and *S. caseolaris*.

|  | *S. alba* |  |  | *S. caseolaris* |  |  |
| --- | --- | --- | --- | --- | --- | --- |
|  | Intergenic | Intron | Coding | Intergenic | Intron | Coding |
| *Copia* | 13.5***(1.43)<sup>†</sup> | 5.8**(0.31) | 4.1***(0.06) | 20.0(0.95) | 8.4(0.09) | 8.0(0.07) |
| *Gypsy* | 38.0(4.12) | 12.4(0.74) | 7.4***(0.18) | 36.7(1.07) | 13.8(0.15) | 15.3(0.14) |
| DNA<br>transposon | 55.8***(0.91) | 23.2**(0.31) | 17.2***(0.23) | 83.1(0.85) | 29.7(0.24) | 30.1(0.26) |
| Others | 112.4***(5.10) | 41.0(1.33) | 14.7***(0.25) | 146.2(4.43) | 41.0(0.81) | 31.6(0.41) |

<sup>†</sup>Upper values in each cell are the number of elements per megabase of DNA in each category and values in parentheses are the proportion of DNA in each location occupied by TEs. Significance levels are shown for comparisons between *S. alba* and *S. caseolaris*, calculated using the Mann-Whitney *U*-test. \*\*, $P < 0.01$ ; \*\*\*, $P < 0.001$ .

Finally, we analyzed the removal rate of LTR elements by estimating the ratio of solo-LTRs to intact elements (S:I). Unequal intra-element homologous recombination resulting in solo-LTRs is known to be one of the major mechanisms eliminating LTR retrotransposons (Devos et al., 2002). As shown in Table 2, the S:I ratio is the lowest for *Gypsy* elements in *S. alba* (0.34), followed by *Copia* in *S. alba* (1.79) and then the *Copia* and *Gypsy* elements in *S. caseolaris* (2.45 and 3.72). Taken together, these results suggest that despite strong purifying selection, high proliferation activity and low elimination rate jointly contributed to the rapid accumulation of *Gypsy* elements in the *S. alba* genome.

**Table 2.** Comparison of the ratios of solo-LTRs to intact elements (S:I) for *Copia* and *Gypsy* LTR retrotransposons between *S. alba* and *S. caseolaris*.

|  | Superfamily | Solo-LTR (S) | Intact Element (I) | S:I |
| --- | --- | --- | --- | --- |
| *S. alba* | *Copia* | 322 | 180 | 1.79 |
|  | *Gypsy* | 264 | 767 | 0.34 |
| *S. caseolaris* | *Copia* | 120 | 49 | 2.45 |
|  | *Gypsy* | 283 | 76 | 3.72 |

### Recent bursts of two *Gypsy* families in *S. alba* are associated with integration site preference

To compare the evolutionary histories of *Copia* and *Gypsy* retrotransposons in the two *Sonneratia* species, we conducted phylogenetic analyses using full length sequences of intact elements for the *Copia* and *Gypsy* superfamilies separately. The maximum-likelihood phylogenetic trees grouped LTR retrotransposon copies of *S. alba* and *S. caseolaris* into 26 *Copia* and 23 *Gypsy* families, half of which (13 out of 26 for *Copia* and 11 out of 23 for *Gypsy*) were shared by both genomes (Supplementary Data: Figure S1). Remarkably, we found that two *Gypsy* families (RLG_1 and RLG_8) are ∼10 times more abundant in *S. alba* (RLG_1: 226 copies and RLG_8: 273 copies) than in *S. caseolaris* (RLG_1: 20 and RLG_8: 22, Supplementary Data: Figure S1). These two *Gypsy* families account for 64.6% of the young copies (< 1 Myr) and 65.1% of the total intact *Gypsy* elements in *S. alba* (Figure 2A). Furthermore, three RLG_1 elements and four RLG_8 elements have identical 5’ and 3’ LTR sequences and perfect 5-bp Target Site Duplications (TSDs), suggesting they are newly originated copies. Therefore, recent bursts of RLG_1 and RLG_8 in *S. alba* are the major contributors to the observed expansion of *Gypsy* elements.

**Figure 2.**
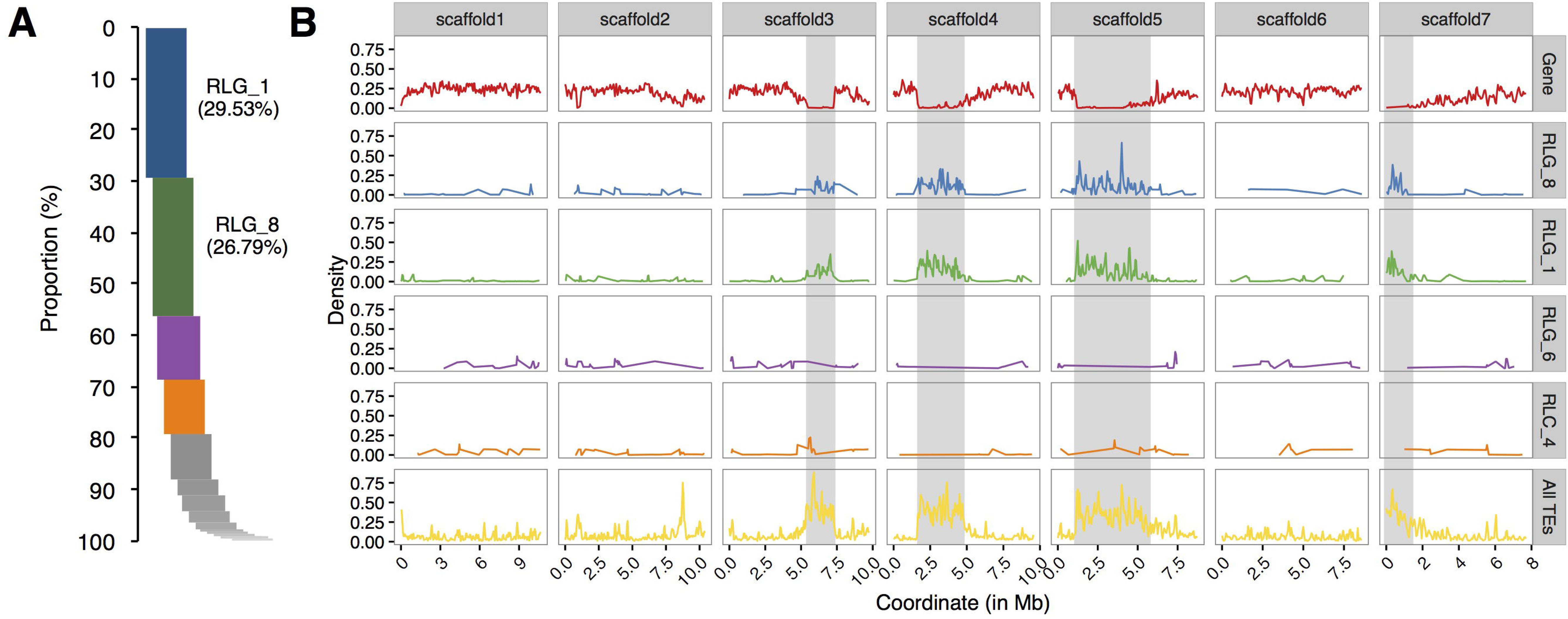
Relative proportion and chromosome distribution of LTR retrotransposon families in *S. alba*. (A) Proportions of intact copy numbers from different LTR retrotransposon families ranked in a descending order. Note that RLG_1 and RLG_8 are the top two abundant LTR retrotransposons in *S. alba*. (B) Chromosome distribution of coding genes (the upper row), the top four abundant LTR retrotransposon families (RLG_1, RLG_8 and RLG_4 of *Gypsy* and RLC_6 of *Copia*), and all TEs along the seven largest scaffolds of the *S. alba* genome. The putative heterochromatin regions (gene density < 0.05) are indicated with the grey shadow. Note that the distributions of RLG_1 and RLG_8 elements track the genomic locations of putative heterochromatic regions but no such pattern is observed for RLG_4 and RLC_6.

Plant LTR retrotransposons are preferentially associated with large heterochromatic regions that flank functional centromeres (Kumar and Bennetzen, 1999). We wondered whether such integration site preference, leading to loose epigenetic repression, might have contributed to the expansion of RLG_1 and RLG_8 families. We scanned scaffolds of the *S. alba* genome using 100 kb sliding windows with a step-size of 50 kb and estimated gene density in each window (see *MATERIALS AND METHODS*). As expected, about half of the RLG_1 copies (50.9%) and 2/5 of the RLG_8 copies (40.7%) are located in gene-poor regions (gene density < 0.05) in *S. alba*. In contrast, the proportion is only 8.1% and 12.5% for the third and fourth most abundant LTR retrotransposon families (Figure 2B).

Further analysis of the structure of RLG_1 and RLG_8 detected a chromodomain (CHD) in the 3’end of both families. CHD was first identified in the heterochromatin protein-1 (HP1) of *Drosophila melanogaster* (Paro and Hogness, 1991) and considered to be involved in chromatin remodeling and the formation of heterochromatin (Paro and Hogness, 1991). Sequence analyses of CHD proteins from diverse organisms classified CHD of the RLG_1 and RLG_8 families into Group I chromodomains (Figure 3A). Both of them possess a highly conservative protein secondary structure (Figure 3B). CHD in *Gypsy* elements, also named chromoviruses, has been experimentally demonstrated to promote the integration of newly emerging retroelements into heterochromatic regions (Gao et al., 2008).

**Figure 3.**
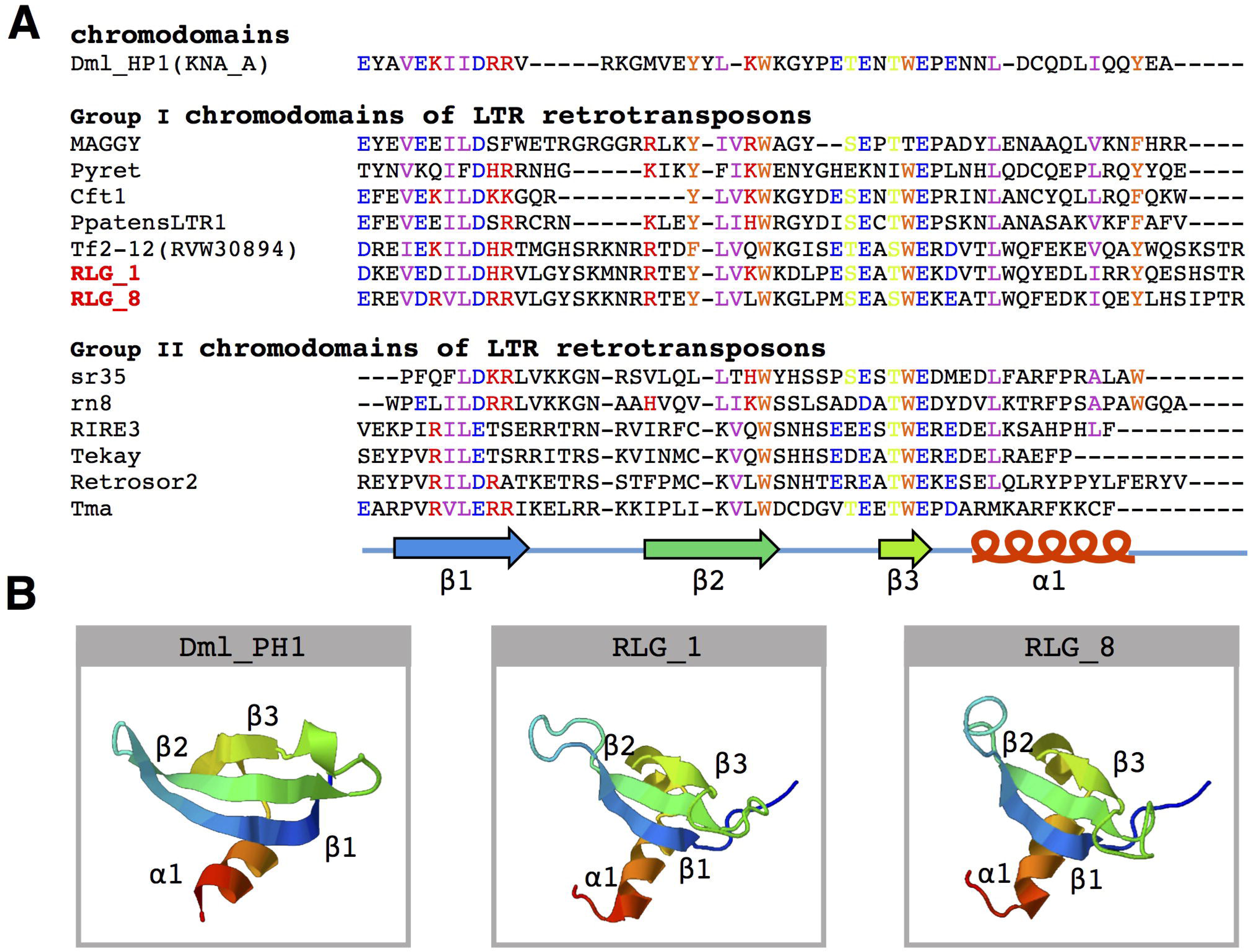
Amino acid sequence alignment and structure prediction of the chromodomain. (A) Multiple alignments of amino acid sequences of known chromodomains and chromo-like motifs from the RLG_1 and RLG_8 families. A schematic representation of β-strands (arrows) and α-helix (cylinder) is on the bottom. The GenBank accession numbers for the sequences used are: DmHP1(KNA_A)-M57574 from *Drosophila melanogaster*; MAGGY-L35053 and Pyret-AB062507 from *Magnaporthe grisea*; Cft1-AF051915 from *Cladosporium fulvum*; PpatensLTR1-XM_001752430 from *Physcomitrella patens*; Tf2-12-RVW30894 from *Vitis vinifera*; sr35-AC068924, rn8-AK068625 and RIRE3-AC119148 from *Oryza sativa*; Tekay-AF050455 from *Zea mays*; RetroSor2-AF061282 from *Sorghum bicolor*; Tma-AF147263 from *Arabidopsis thaliana*. (B) Predicted three-dimensional structure of chromodomains from the DmHP1 protein (M57574) and the RLG_1 and RLG_8 families.

### Expanded *Gypsy* families are under relaxed epigenetic repression

If CHD indeed promotes the insertions of RLG_1 and RLG_8 into heterochromatin regions, we would expect that epigenetic repression of these elements is relaxed in the host genome. In plants, CHH methylation mediated by 24-nt siRNAs is responsible for the establishment of *de novo* methylation against the emergence of new TE copies (Law and Jacobsen, 2010). We thus estimated correlations between gene density, CHH methylation level, and 24-nt siRNA abundance for the RLG_1 and RLG_8 copies in *S. alba*. Gene density of the window in which each RLG_1 and RLG_8 copy resides was used in this analysis.

For both RLG_1 and RLG_8, copies residing in the gene-poor regions (gene density < 0.05) exhibited a significantly lower level of CHH methylation (RLG_1: median, hereafter, 10.6% and RLG_8: 15.9%) than the rest (RLG_1: 44.0% and RLG_8: 37.7%, Mann-Whitney *U*-test, all P < 0.001, Figure 4A). The 24-nt siRNA targeting pattern is in accordance with the difference in methylation levels, i.e., a lower level of 24-nt siRNA abundance on TEs residing in gene-poor regions than the other TEs (4.0 vs. 7.5 reads per element per kilobase for RLG_1 and 2.9 vs. 5.8 for RLG_8, Mann-Whitney *U*-test, all P < 0.001; Figure 4A). These results support the idea that members of RLG_1 and RLG_8 have experienced relaxed epigenetic repression associated with their insertion preference. Gene-poor regions might represent a “safe haven” for these elements to coexist with host cells.

**Figure 4.**
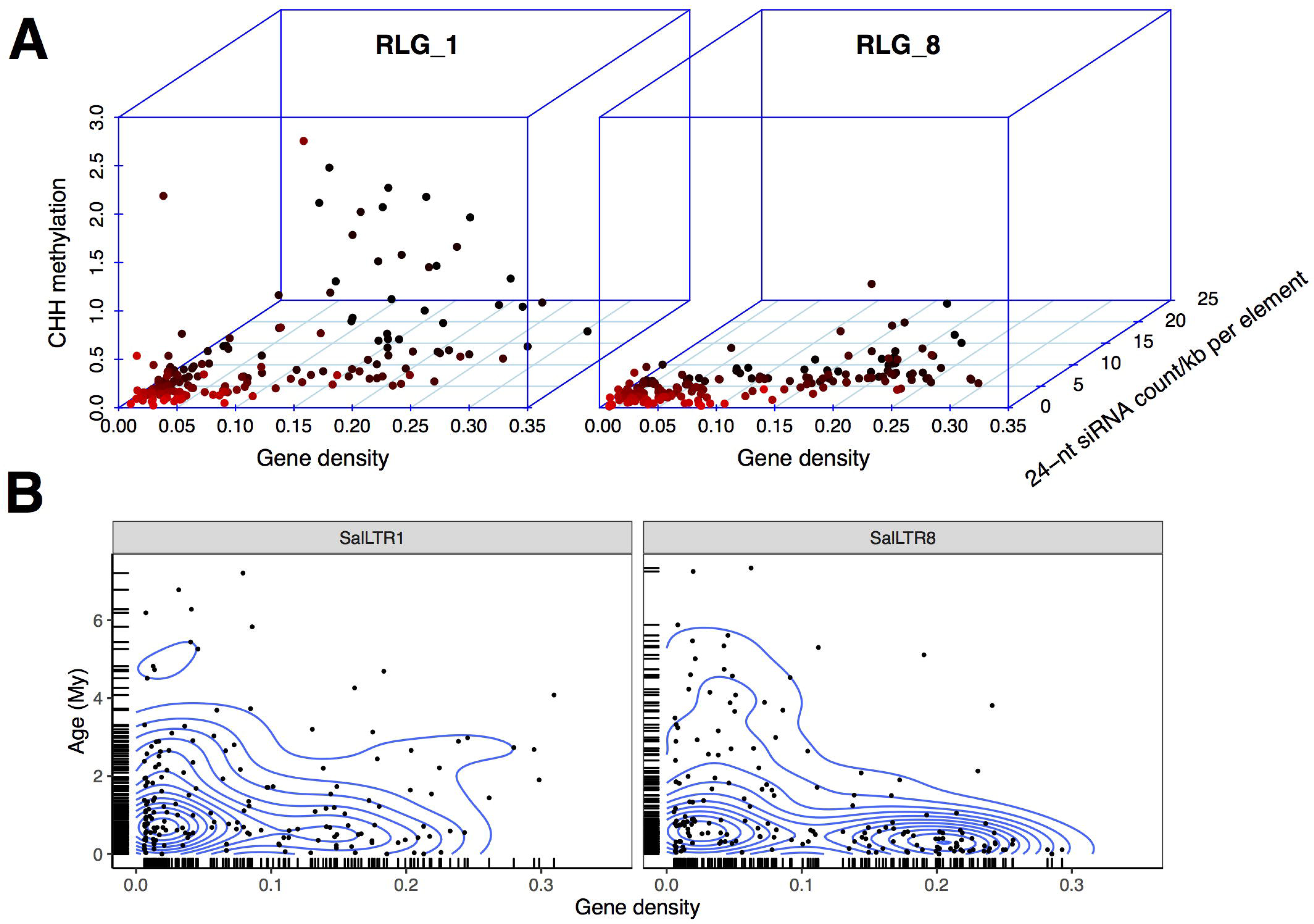
Epigenetic repression and insertional locations of RLG_1 and RLG_8 in *S. alba*. (A) Correlations between CHH methylation level, 24-nt siRNA abundance of the RLG_1 and RLG_8 elements, and gene density of the 100 Kb region in which each retroelement is located in the *S. alba* genome. Each dot represents an intact RLG_1 or RLG_8 element and is colored in red to black gradient according to gene density of the genomic region in which the element is located. (B) Age distribution of the RLG_1 or RLG_8 elements along genomic regions with different gene density. Contour lines represent estimated two dimensional kernel density.

### Weak effect of TE expansion on salt stress gene expression in *S. alba*

Despite the overall preference for integration into gene-poor regions, there is still a considerable fraction of young copies (< 1 Myr) of RLG_1 (21.8%) and RLG_8 (45.2%) inserted into genomic regions with gene density > 0.15 (Figure 4B). It is possible that recently amplified TEs can affect nearby gene expression by carrying repressive epigenetic markers or creating new regulatory elements (Naito et al., 2009, Chuong et al., 2017). Recently, Feng et al. (2020) reported a comparative transcriptome study of *S. alba* treated with 0, 250mM, and 500mM NaCl. They used the 250mM NaCl condition as a control and 0 and 500mM NaCl as stresses with low or high salt (Feng et al., 2020). Using these data, we asked whether intact LTR retrotransposons in *S. alba* can be activated by salt stress and how TE expansion might affect gene expression under stress conditions.

Many more of the 767 and 180 intact *Gypsy* and *Copia* retrotransposons were expressed in roots (*Gypsy*: 420 and *Copia*: 15) than in leaves (*Gypsy*: 169 and *Copia*: 11) (Figure 5A). This is coincident with the fact that roots are directly exposed to the saline environment. An intact retrotransposon was considered to be expressed if it had more than 5 reads in both biological replicates under at least one of the low, medium, or high salinity conditions. There was more upregulation (log2 fold change > 1) than downregulation (log2 fold change > -1) for these retroelements in all comparisons, most prominently for the comparison between 500mM and 250mM NaCl in roots (Figure 5A). However, only four retrotransposons were significantly mis-regulated due to large variation among replicates.

**Figure 5.**
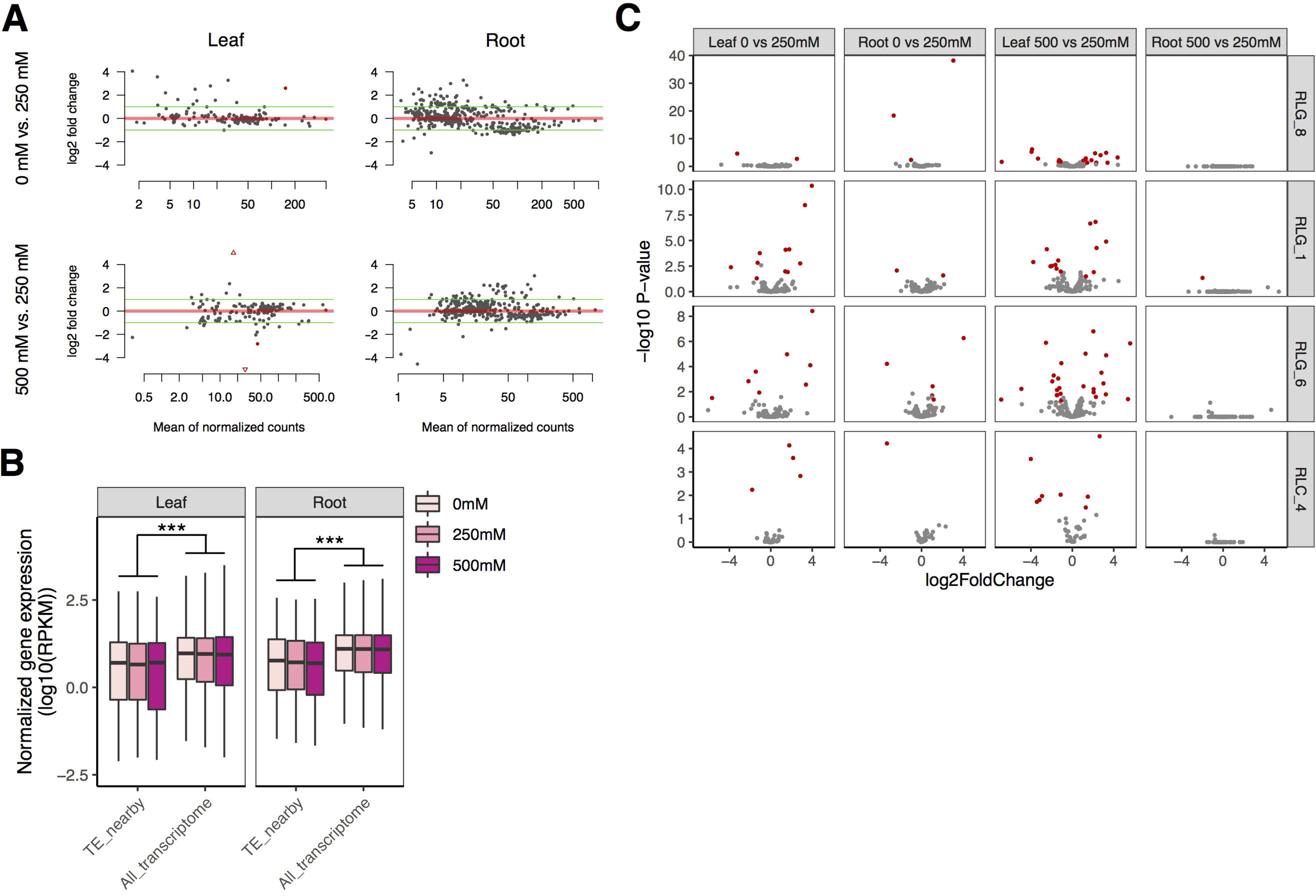
Expression patterns of TEs and their flanking genes under salt treatments in *S. alba*. (A) Fold changes of the expression levels of intact *Gypsy* and *Copia* retrotransposons across salt treatments (the upper: 0 mM vs 250 mM NaCl, the lower: 250 mM vs 500 mM NaCl). (B) Expression levels of genes neighboring copies of the four most abundant LTR retrotransposon families in comparison with the whole transcriptome. (C) Fold expression change of genes next to copies of the four most abundant LTR retrotransposon families across salt treatments (the upper: 0 mM vs 250 mM NaCl, the lower: 500 mM vs 250 mM NaCl). Genes with significant mis-expression (FDR < 0.05 and ≥ 2-fold change) between treatments are indicated in red.

We further checked changes of expression in genes with intact TEs in introns, within 3kb upstream of the transcription start site, or downstream of the transcription termination site for the top four most abundant LTR retrotransposon families in *S. alba*. Gene expression level was calculated as reads per kb per million reads (RPKM). For all the four families, genes with neighboring TEs were expressed at a significantly lower level than the whole transcriptome under all conditions (Mann-Whitney *U*-test, all P < 0.001 in both leaves and roots; Figure 5B), supporting the view that TEs carry repressive epigenetic marks that may reduce flanking gene expression (Naito et al., 2009, Chuong et al., 2017). While a few TE-adjacent genes were significantly mis-regulated (FDR < 0.05 and fold change ≥ 2) by salt (Figure 5C and Supplementary Data: Table S2), this set of loci was not enriched in mis-expressed genes relative to the whole transcriptome in any of our comparisons (Fisher exact test, all P > 0.05, Supplementary Data: Table S3). We also estimated correlations between TE expression levels and their nearby genes in leaves or roots across the three salt conditions (Supplementary Data: Table S4). Only two significantly correlated pairs were identified in leaves: a RLC_4 copy and its 1307-bp upstream gene of unknown function (Pearson’s correlation, *r* = 0.99, adjusted P < 0.05), and a RLG_8 copy and its 505-bp downstream gene *Abscisic acid 8’-hydroxylase 3* (Pearson’s correlation, *r* = 0.99, adjusted P < 0.05). *Abscisic acid 8’-hydroxylase* catalyzes the first step in the oxidative degradation of (+)-ABA (Krochko et al., 1998). Among the four ABA 8’-hydroxylase genes in *Arabidopsis*, *CYP707A3* supports the drought tolerance of plants via maintenance of ABA levels (Umezawa et al., 2006).

Besides intact LTR retrotransposons, truncated elements and solo-LTRs may also affect expression of nearby genes. We then examined expression changes of all the truncated retrotransposons (3,279) and solo-LTRs (586) as well as expression changes of their nearby genes in *S. alba* as described above. Similar with the intact elements, the truncated LTR retrotransposons were expressed more in roots (615) than in leaves (355), and exhibited more upregulation (log2 fold change > 1) than downregulation (log2 fold change > -1) upon high salinity treatment (Supplementary Data: Figure S2). In contrast, solo-LTRs were rarely expressed (Supplementary Data: Figure S2). A total of 24 truncated retrotransposons and one solo-LTR were significantly mis-expressed in roots or leaves between treatments. Two TE-gene pairs were found to have significant expression correlation across conditions of different salt treatments. One pair was identified in leaves, involving a truncated RLG_6 copy (RLG_6_T1) and its 1306-bp downstream gene *thylakoid soluble phosphoprotein of 9 kDa (TSP9)* (Pearson’s correlation, *r* = 0.99, adjusted P < 0.01). TSP9, is a plant-specific protein in the photosynthetic thylakoid membrane that regulates light harvest in *Arabidopsis* (Carlberg et al., 2003; Fristedt et al., 2009). The other pair was identified in roots, involving another truncated RLG_6 copy (RLG_6_T2) and its 705-bp upstream gene *glutamate-tRNA ligase,* also known as *glutamyl-tRNA synthetase* (Pearson’s correlation, *r* = -0.99, adjusted P < 0.05).

Taken together, we identified four TE-gene pairs with significant expression correlation in *S. alba* that involve both intact and truncated LTR retrotransposons. Among the *S. caseolaris* orthologs of these four genes, only *TSP9* had a truncate RLG_6 copy inserted in its downstream region. However, none of the RLG_6 copies was expressed in leaves of *S. caceolaris* under normal condition (Wang et al., unpublished data). Orthologs of the other three genes in *S. caseolaris* had no insertion of RLC_4, RLG_8 or RLG_6 (BLASTn, all *e*-value > 0.05) in their flanking 10-kb up- or downstream region.

### Potential *cis*- and *trans*-acting roles of TEs on gene expression in *S. alba*

It is very well known that LTRs contain CREs that can confer stress-inducible expression pattern on TEs and their flanking genes (Cavrak et al., 2014). To infer the potential *cis*-acting role of LTR retrotransposons, we predicted CREs in LTRs for each family using PlantCARE (Lescot et al., 2002). Consensus sequences of *Gypsy* and *Copia* families that have more than ten copies were used in this analysis. We identified abundant CREs responsive to various biotic and abiotic stresses in LTRs, whereas none of these CREs is responsive to salt stress (Figure 6). RLG_1 and RLG_8 contained CREs responsive to anoxic, drought, low temperature, pathogen and fungal elicitor, which are not more abundant than CREs residing in LTRs of other families (Figure 6). Interestingly, the LTRs of RLG_8 harboring three ABA-responsive elements (ABREs) involved in ABA and drought responsiveness (Figure 6) is coincident with a RLG_8 copy positively co-expressed with *Abscisic acid 8’-hydroxylase 3* the gene nearby (see above). ABREs and MYB binding sites (MBSs) involved in drought inducibility were also found in the other three retrotransposons that showed significant expression correlation with their flanking genes (Figure 6). These results suggest that *cis*-regulatory role of LTR retrotransposons in *S. alba,* although rare, might occur via cross talk between drought and salt stress signaling pathways.

**Figure 6.**
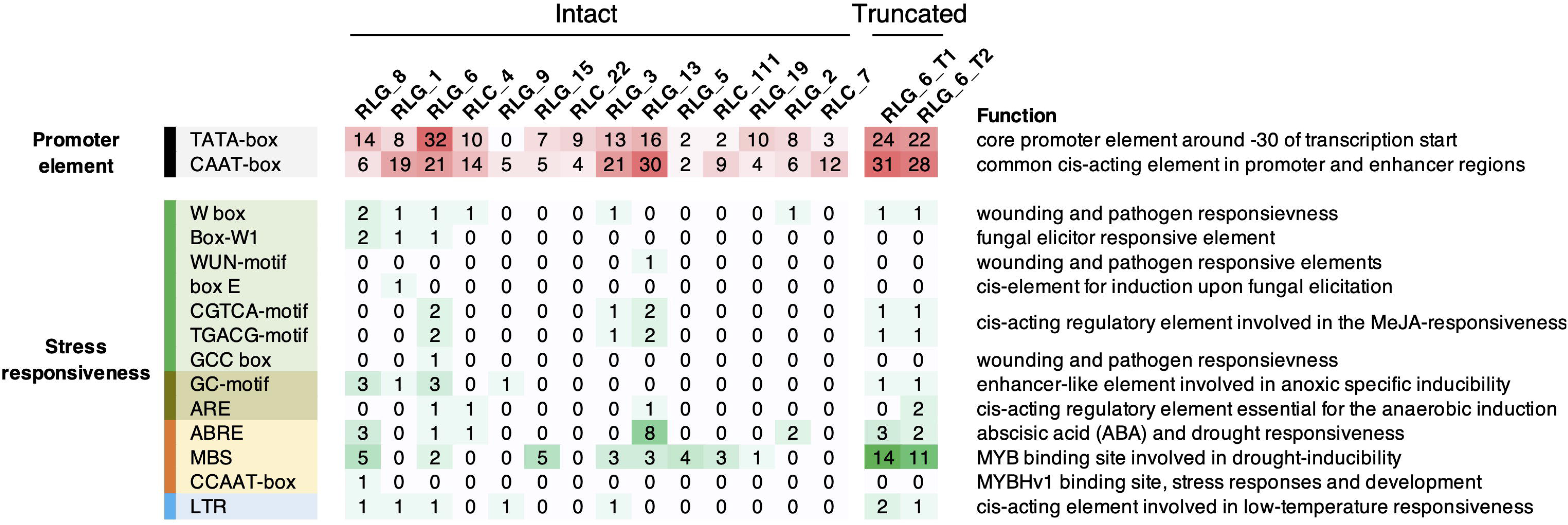
Summary of *cis*-acting regulatory elements (CREs) in the LTRs of *Gypsy* and *Copia* families with more than ten copies in *S. alba*. Also shown are CREs in the full length of the two truncated elements (RLG_6_T1 and RLG_6_T2) which significantly correlated with their nearby genes in expression levels. Color depths and numbers indicate the abundance of specific CREs.

Besides *cis*-acting regulation, TEs can also function in *trans,* either resembling microRNA (miRNA) genes (Li et al., 2011) or acing as miRNA sponges competing with target transcripts of the same miRNA (Cho and Paszkowski, 2017; Cho, 2018; Palazzo and Koonin, 2020). To test these possibilities in *S. alba*, we annotated known and novel miRNAs using small RNA sequencing data described above with miRDeep2 (Friedländer et al., 2012) and predicted miRNA targets using psRNATarget (Dai and Zhao, 2011). An expectation score of 2.5 was used as the cutoff in the target prediction. All LTR retrotransposons including the intact and truncated elements and solo-LTRs were used in these analyses. We identified 105 known and 14 novel miRNAs expressed (Reads per million mapped reads, RPM > 1) in both biological replicates; none was derived from TEs (Supplementary Data: Table S5). We then used the criteria described by Cho and Paszkowski (2017) to identify TE-gene pairs with potential miRNA competition. Among the 65 LTR retrotransposons predicted to contain potential miRNA binding sites, only 21 were expressed in roots or leaves of *S. alba* under either condition of salt treatments (Supplementary Data: Table S6). Requiring co-expression of both TEs and genes, we identified six TE-gene pairs with sequence matching in sense orientation and miRNA-binding sites within the matching regions. Nevertheless, none of them showed significantly expression correlation across different salt treatment conditions (Pearson correlation, all P > 0.05, Supplementary Data: Table S7). Therefore, *trans*-acting role of LTR retrotransposons is negligible in *S. alba*.

## DISCUSSION

In her seminal studies of maize, McClintock proposed that TEs are “controlling elements” that may reprogram genome responses to environmental challenges (McClintock, 1984). Although still controversial, this idea has been supported by increasing empirical evidence (Naito et al., 2009, Cavrak et al., 2014, Horvath et al., 2017). TEs can be activated by stress, generate genetic variation, and modulate gene expression (Feschotte, 2008, Henaff et al., 2014). While induced TE activity has long been thought to benefit the host, it is puzzling that mangroves have experienced convergent reduction of TE loads despite being adapted to extreme environments with limited standing genetic variation (Guo et al., 2018, Lyu et al., 2018). How often TE activities are regulated by environmental cues and how these activities may contribute to a significant adaptive genome response in natural environments remain intriguing questions.

In asking whether divergent habitats promote differential TE accumulation, we found TEs that were accumulating more in *S. alba* than in *S. caseolaris* despite the overall tendency of unloading the TE burden in mangroves (Figure 1A,B). This result, together with the fact that *S. alba* faces higher salinity than *S. caseolaris,* is consistent with the view that TEs tend to be active in plants growing in challenging habitats. The large fraction of young copies in *S. alba* (Figure 1C,D) suggests that TE originations are more frequent in *S. alba* than *S. caseolaris.* As young elements do not have enough time to degenerate, genome purging mechanisms should have little effect, if any, on their copy number. Although we do not have direct evidence that increased mobility in *S. alba* is triggered by environmental stress, transcriptome analyses suggest that more TEs are constitutively expressed in roots than leaves (Figure 5A), supporting this idea. Interestingly, purifying selection against TE insertions was even stronger in *S. alba* than in *S. caseolaris* (Table 1). The high origination rate and strong purifying selection might have led to fast turnover of TEs in *S. alba*, echoing a previous report of constant TE-host conflict masked by unloading of the TE burden in *R. apiculata* (Wang et al., 2018). Difference in unequal homologous recombination, as inferred by the S:I ratio, (Table 2) may further contribute to the differential TE accumulation between the two *Sonneratia* species.

The recent burst of TE abundance is particularly prominent for the *Gypsy* retrotransposons in *S. alba*. We found that the two most abundant *Gypsy* families in *S. alba* (RLG_1 and RLG_8) mainly accumulate in gene-poor regions. This is likely due to a balance between TE integration site preference and post-integration selection processes (Sultana et al., 2017). The presence of the CHD domain in RLG_1 and RLG_8 likely drive heterochromatic integration site preference. However, the same integration site preference should be maintained in *S. caseolaris* if the CHD domain is sufficient to locate TE insertions to the same genomic region. Therefore, integration site preference alone cannot explain the difference in TE originations between species. Alternatively, if the two families differed in integration site preference between species, e.g. because of divergence of host proteins involved in TE tethering, the amplified copies should be much older than our observation (< 1 Myr), given that *S. alba* and *S. caseolaris* have diverged for ∼11.6 Myr. We propose that the differential accumulation of RLG_1 and RLG_8 between species was probably triggered by environmental stress. It is possible that severe environments reactivated TEs more in *S. alba* than in *S. caseolaris* and then heterochromatin provided a “safe haven” for the accidentally inserted copies, allowing them to proliferate rapidly with relaxed epigenetic repression. The synergy between these mechanisms may constitute a self-reinforcing and self-perpetuating cycle of TE origination leading to long-term maintenance of genetic diversity. This hypothesis needs to be rigorously tested using data on stress-activated retrotransposition. We do see more up- than down-regulation of RLG_1 and RLG_8 during salt stress (Supplementary Data: Figure S3), suggesting stress can activate TE transcription. The presence of CREs responsive to anoxia and drought but not to salt in the LTRs of RLG_1 and RLG_8 suggests that the cross-talk between different stress signaling pathways might play a significant role in the potential TE activation in *S. alba*.

Although stress-induced TE activity in *S. alba* is plausible, our transcriptome analyses suggest that TE expansion into regulatory sequences is rare. This is evident in the observation that TEs are mainly expressed in roots whereas the vast majority of differentially expressed TE-adjacent genes were found in leaves (Figure 5). Comparative transcriptome analyses between *S. alba* and *S. caceolaris* also suggest that TE expansion in *S. alba* influences between-species expression divergence mainly via carrying repressive epigenetic markers rather than creating new regulatory elements (Supplementary Note). There are only four potential cases of TE co-option in *S. alba* (Supplementary Data: Table S4). In one case, unlike constitutive expression of *CYP707A3* in *Arabidopsis*, *Abscisic acid 8’-hydroxylase 3* in *S. alba* and the RLG_8 copy upstream of it exhibited salt-induced expression in leaves only under 500mM NaCl, leading to a significant positive correlation in their expression levels (Supplementary Data: Table S4). This result together with the presence of ABREs in the LTRs of RLG_8 suggests that the RLG_8 copy might have been integrated into a stress-responsive gene regulatory network involving ABA catabolism. In *Arabidopsis, cyp707a3* mutant plants exhibit enhanced drought tolerance (Umezawa et al., 2006). Our result might also suggest that the strong salt tolerance of *S. alba* is achieved via reduced ABA degradation. We were unable to make any biological inferences in the other case of the RLC_4 copy, since the gene next to it has no known function. The rest two cases both involved truncated RLG_6 copies that possess tens of MBSs involved in drought-inducibility (Figure 6). The positive expression correlation of one RLG_6 copy with *TSP9* in leaves and the negative expression correlation of another RLG_6 copy with *glutamate-tRNA ligase* in roots suggest that TEs might mediate the cross-talk between photosynthesis and nitrogen metabolism during the adaptation of *S. alba* to mangrove swamps.

It should be noted that a retrotransposon is potentially responsive to diverse stress conditions. As we only examined one stress condition (salinity), it is possible that co-option of TEs under other stress conditions was missed in this study. Nevertheless, it has been shown that enhancers often evolve from ancient TE sequences (Lynch et al., 2015, Villar et al., 2015). The evolutionary age of a TE was found to correlate positively with its likelihood of contributing to a regulatory element in human and mouse (Simonti et al., 2017). As the vast majority of retrotransposons in *S. alba* are relatively young, they may not have had enough time to get integrated into gene regulatory networks unless strong selection favors maintenance of the TE insertions, such as the *Hopscotch* insertion increasing *tb1* expression in maize (Studer et al., 2011). Considering the small effective population size of *S. alba* (Zhou et al., 2007, Zhou et al., 2011), natural selection may not be efficient enough to drive rapid fixation of TE insertions even though some TEs are capable of modulating expression of nearby genes.

In summary, our results together with a previous study (Wang et al., 2018) suggest that expansion of *Gypsy* retrotransposons, probably triggered by stress, is prevalent in mangroves despite the overall tendency of unloading the TE burden. The extent of TE proliferation across families is influenced by integration site preference and is subject to epigenetic repression and genome purging mechanisms. Although genetic variation generated by TE activities may be beneficial to host adaption, TE exaptation in *S. alba* is rare at least in terms of salt stress. Further experiments are needed to test the hypothesis of rapid transcriptional regulatory rewiring mediated by TE mobilization even though this hypothesis is appealing for extremophiles in which stress-induced reactivation of TEs is thought to be frequent.

## [Supplementary Information]

**Figure S1.** Maximum likelihood (ML) phylogenetic trees of *Copia* and *Gypsy* LTR retrotransposon families.

**Figure S2.** Expression fold changes of the truncated LTR retrotransposons and solo-LTRs as well as their nearest genes across salt treatments in leaves and roots. TEs or genes with significant mis-expression (Benjamini-Hochberg FDR < 0.05) between treatments are indicated in red. Green lines indicate two-fold expression changes.

**Figure S3.** Expression level fold change of intact RLG_1 and RLG_8 retrotransposons across salt treatments.

**Table S1.** List of all intact LTR retrotransposons in *Sonneratia alba* and *S. caseolaris.* (Excel)

**Table S2.** TE-adjacent genes in *Sonneratia alba* that were significantly mis-regulated by salt treatment. (Excel)

**Table S3.** The number of TE-adjacent genes in *Sonneratia alba* that were significantly mis-regulated by salt treatment.

**Table S4.** Correlations between expression levels of TEs and their nearby genes in *Sonneratia alba*. (Excel)

**Table S5.** Annotation of known and new miRNAs in leaves of *S. alba*. (Excel)

**Table S6.** LTR retrotransposons predicted to be miRNA targets in *S. alba*. (Excel)

**Table S7.** Pairs of LTR retrotransposons and genes with potential miRNA competition in *S. alba.* Pearson correlation was calculated for expression levels of each pair across 0mM, 250 mM and 500mM salt treatments in roots and leaves, respectively. *cor*, Pearson correlation coefficient; P, P value.

