## Supplementary material for "Weak effect of *Gypsy* retrotransposon bursts on *Sonneratia alba* salt stress gene expression": Figure S1-S3, Table S3, Table S7

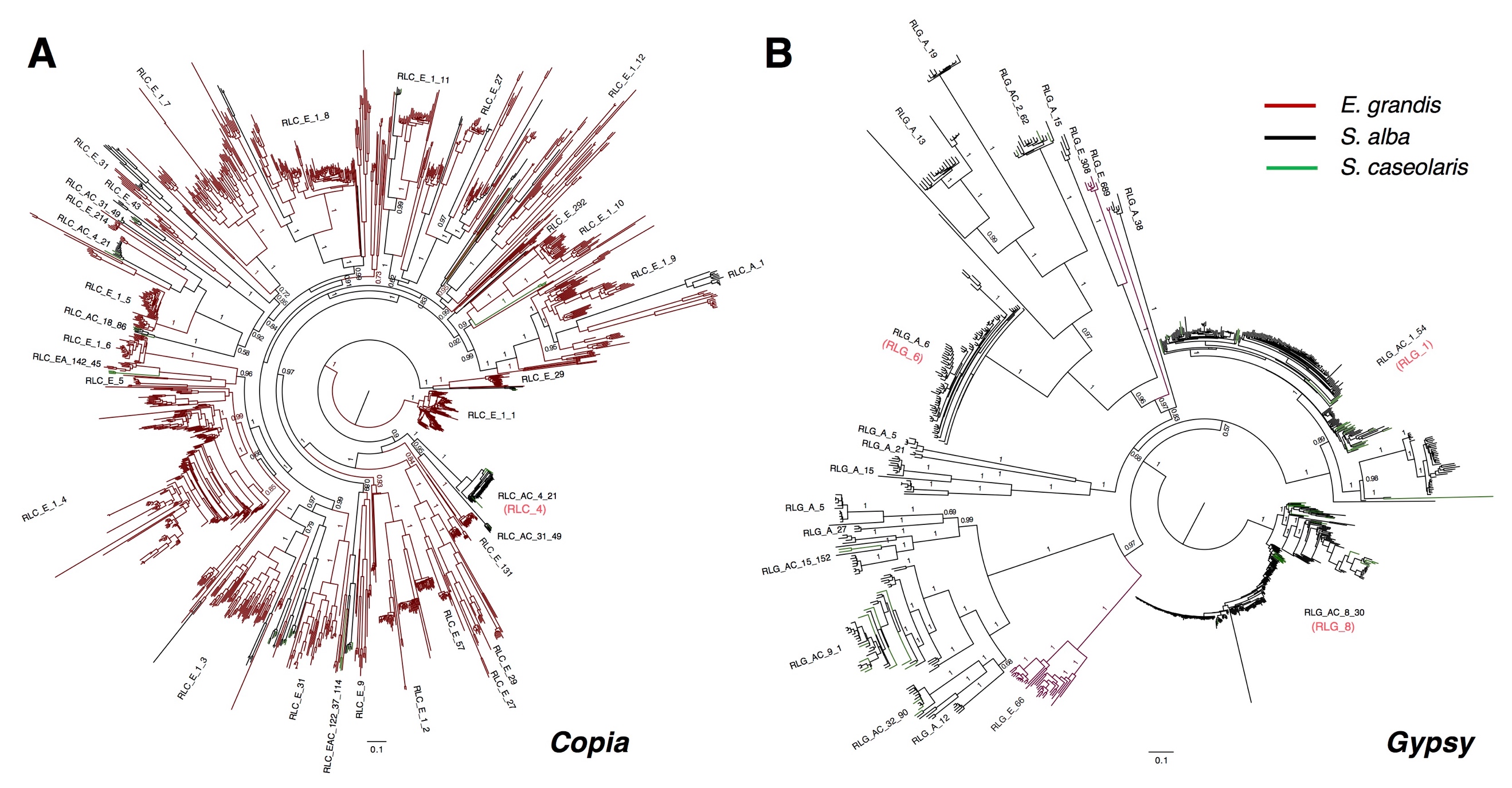

**Figure S1** Maximum likelihood (ML) phylogenetic trees of *Copia* and *Gypsy* LTR retrotransposon families. Sequences of intact (A) *Copia* and (B) *Gypsy* copies identified in *S. alba* and *S. caseolaris* were used for this analysis. Supporting values of each branch were calculated by resampling estimated log-likelihoods (RELL)-like local support values.

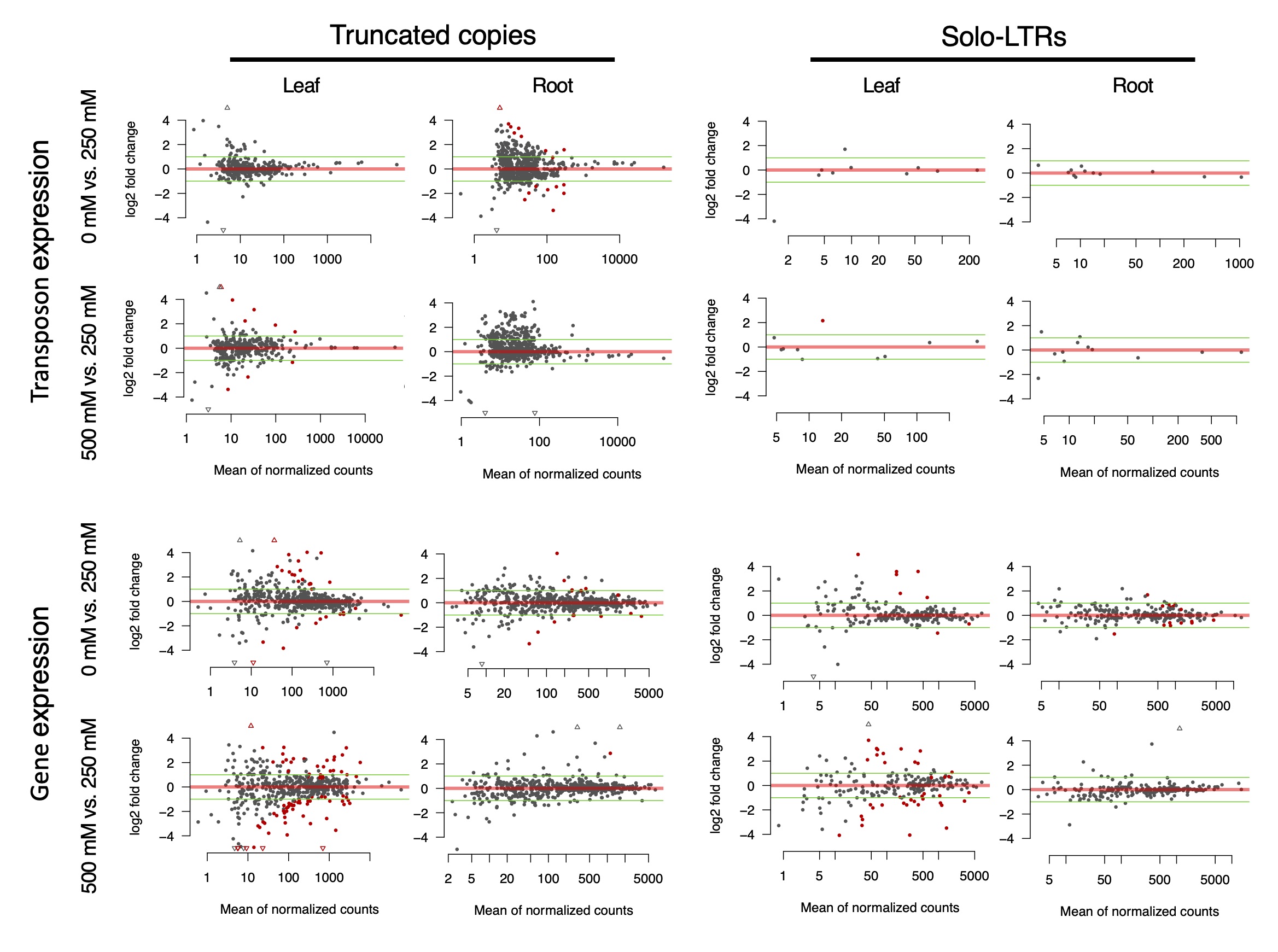

**Figure S2** Expression fold changes of the truncated LTR retrotransposons and solo-LTRs as well as their nearest genes across salt treatments in leaves and roots. TEs or genes with significant mis-expression (Benjamini-Hochberg FDR < 0.05) between treatments are indicated in red. Green lines indicate two-fold expression changes.

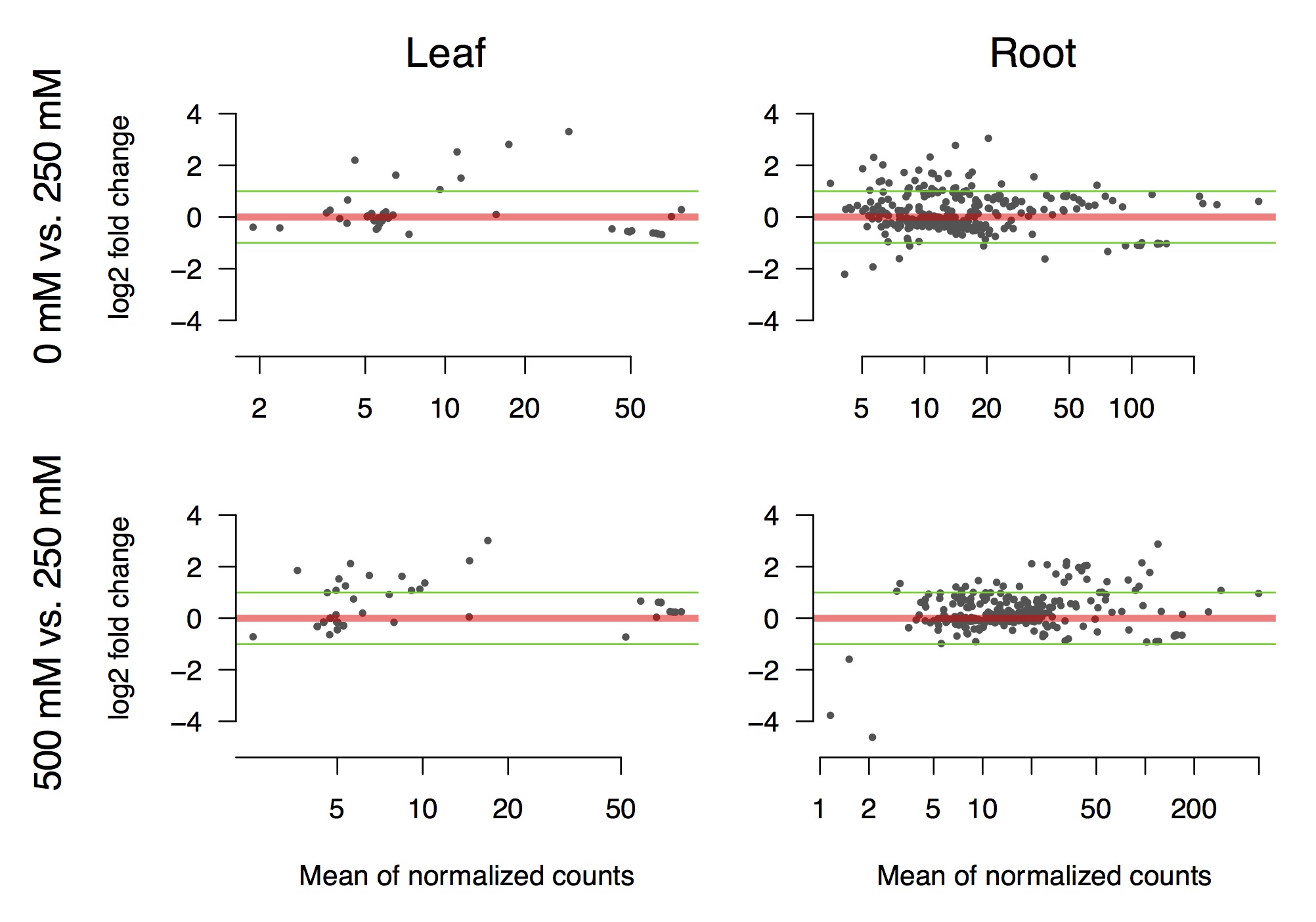

**Figure S3** Expression level fold change of intact RLG_1 and RLG_8 retrotransposons across salt treatments. Upper panels: 0 mM vs 250 mM NaCl, lower panels: 250 mM vs 500 mM NaCl. Transposon elements with significant mis-expression (Benjamini-Hochberg FDR < 0.05) between treatments are indicated in red.

**Table S1**. List of all intact LTR retrotransposons in *Sonneratia alba* and *S. caseolaris.* (Excel)

**Table S2**. TE-adjacent genes in *Sonneratia alba* that were significantly mis-regulated by salt treatment. Only those from of the four most abundant LTR retrotransposon families were used in this analysis. (Excel)

**Table S3**. The number of TE-adjacent genes in *Sonneratia alba* that were significantly mis-regulated by salt treatment. Only those from the top four most abundant LTR retrotransposon families were counted.

|  |  | Leaf 0 vs. 250 mM | Root 0 vs. 250 mM | Leaf 500 vs. 250 mM | Root 500 vs. 250 mM |
| --- | --- | --- | --- | --- | --- |
| RLG_8 | Differentially expressed | 2 | 3 | 18 | 0 |
|  | Total expressed | 108 | 99 | 106 | 126 |
| RLG_1 | Differentially expressed | 11 | 2 | 14 | 1 |
|  | Total expressed | 165 | 159 | 155 | 193 |
| RLG_6 | Differentially expressed | 8 | 5 | 25 | 0 |
|  | Total expressed | 162 | 145 | 162 | 203 |
| RLC_4 | Differentially expressed | 4 | 1 | 8 | 0 |
|  | Total expressed | 46 | 41 | 44 | 57 |
| Whole transcriptome | Differentially expressed | 1,212 | 390 | 2,835 | 59 |
|  | Total expressed | 26,326 | 27,243 | 26,326 | 27,243 |

**Table S4**. Correlations between expression levels of TEs and their nearby genes in *Sonneratia alba*. Expression levels in leaves or roots across three salt conditions were used for this analysis. (Excel)

**Table S5**. Annotation of known and new miRNAs in leaves of *S. alba*. (Excel)

**Table S6**. LTR retrotransposons predicted to be miRNA targets in *S. alba*. (Excel)

**Table S7**. Pairs of LTR retrotransposons and genes with potential miRNA competition in *S. alba.* Pearson correlation was calculated for expression levels of each pair across 0mM, 250 mM and 500mM salt treatments in roots and leaves, respectively. *cor*, Pearson correlation coefficient; P, P value.

| LTR elements | Shared miRNA | miRNA sequence | Gene | Leaf | |  | Root | |
| --- | --- | --- | --- | --- | --- | --- | --- | --- |
|  |  |  |  | *cor* | P |  | *cor* | P |
| SalLTR13_scaffold1_7261969_7268972 | miR3937 | acaggcgguggaucaaauaugaau | evm.model.scaffold11.642 | -0.14 | 0.79 |  | 0.60 | 0.21 |
| SalLTR6_scaffold50_292888_294854 | miR5622 | auuagcuguugggacuuaaaagcc | evm.model.scaffold39.173 | -0.74 | 0.09 |  | 0.78 | 0.07 |
|  |  |  | evm.model.scaffold12.225 | -0.44 | 0.39 |  | -0.02 | 0.97 |
| SalLTR6_scaffold50_831587_837043 | miR4239 | auuguuauuuuguuggaccggccu | evm.model.scaffold14.309 | -0.37 | 0.47 |  | 0.06 | 0.91 |
| SalLTR6_scaffold50_881694_887933 |  |  | evm.model.scaffold3.1127 | -0.61 | 0.20 |  | -0.07 | 0.90 |
| SalLTR6_scaffold50_878576_885125 |  |  | evm.model.scaffold22.100 | -0.31 | 0.55 |  | -0.08 | 0.88 |
| SalLTR1_scaffold5_755227_760280 | miR8051 | uauuucuuggacucggcuuguaac | evm.model.scaffold54.63 | -0.12 | 0.82 |  | -0.02 | 0.97 |
